# *In vitro* and *in vivo* interaction of caspofungin with isavuconazole against *Candida auris* planktonic cells and biofilms

**DOI:** 10.1101/2021.03.08.434267

**Authors:** Fruzsina Nagy, Zoltán Tóth, Fanni Nyikos, Lajos Forgács, Ágnes Jakab, Andrew M. Borman, László Majoros, Renátó Kovács

**Author notes:** Corresponding author: Renátó Kovács; Department of Medical Microbiology, Faculty of Medicine, University of Debrecen, 4032 Debrecen, Nagyerdei krt. 98., Hungary.

## Abstract

The *in vitro* and *in vivo* efficacy of caspofungin was determined in combination with isavuconazole against *Candida auris*. Drug–drug interactions were assessed utilising the fractional inhibitory concentration indices (FICIs), the Bliss independence model and an immunocompromised mouse model. Median planktonic minimum inhibitory concentrations (pMICs) of 23 *C. auris* isolates were between 0.5 and 2 mg/L and between 0.015 and 4 mg/L for caspofungin and isavuconazole, respectively. Median pMICs for caspofungin and isavuconazole in combination showed 2–128-fold and 2–256-fold decreases, respectively. Caspofungin and isavuconazole showed synergism in 14 out of 23 planktonic isolates (FICI range 0.03–0.5; Bliss cumulative synergy volume range 0–4.83). Median sessile MICs (sMIC) of 14 biofilm-forming isolates were between 32 and >32 mg/L and between 0.5 and >2 mg/L for caspofungin and isavuconazole, respectively. Median sMICs for caspofungin and isavuconazole in combination showed 0–128-fold and 0-512-fold decreases, respectively. Caspofungin and isavuconazole showed synergistic interaction in 12 out of 14 sessile isolates (FICI range 0.023–0.5; Bliss cumulative synergy volume range 0.13–234.32). In line with the *in vitro* findings, synergistic interactions were confirmed by *in vivo* experiments. The fungal kidney burden decreases were more than 3 log volumes in mice treated with combination of 1 mg/kg caspofungin and 20 mg/kg isavuconazole daily; this difference was statistically significant compared with control mice (p<0.001). Despite the favourable effect of isavuconazole in combination with caspofungin, further studies are needed to confirm the therapeutic advantage of this combination when treating an infection caused by *C. auris*.

## 1. Introduction

Since its first identification more than 10 years ago, *Candida auris* has emerged as a global public health threat due to its ability to cause nosocomial outbreaks of invasive infections in health care facilities worldwide [1]. Previously, four major phylogenetically distinct lineages (South Asian, East Asian, South African and South American) emerged simultaneously, a phenomenon that highlights the global dissemination of this pathogen. In addition, a potential fifth clade (Iranian origin) has also been described in the recent past [2–3].

*C. auris* can colonise a variety of body sites and medical implants such as central venous catheters, where biofilm development is one of the most important complications [4]. Clinical studies have shown that indwelling devices were the source in 89% of *C. auris* bloodstream infections; these data emphasise the clinical importance of these sessile communities [5–6]. It is clear that *C. auris* has exceptionally high minimum inhibitory concentrations (MICs) against the three main classes of antifungals [7–9]; therefore, the potential biofilm-forming ability further complicates treatment [10]. For example, echinocandins – including caspofungin – are frequently administered for the treatment of invasive *C. auris* infections [11–12]. However, these drugs are not expected to be effective in biofilm-related *C. auris* diseases due to the 2–512-fold higher sessile MIC values to echinocandins [6]. The need for novel therapeutic approaches against *C. auris* is increasing, but the development of new antifungal drugs has decelerated. Therefore, a promising treatment strategy would be to administer antifungals in combination, an approach that can reduce the toxicity and improve the pharmacokinetics and the antifungal effect of drugs used, ultimately improving the prognosis of patients [13, 14].

In 2016, a new broad-spectrum antifungal drug, isavuconazole, was introduced in clinical practice; it has a favourable safety profile with high activity against a wide variety of fungal pathogens, but the activity of isavuconazole against *C. auris* is variable [15]. Nevertheless, a multicenter study revealed that isavuconazole was not inferior relative to caspofungin for the primary treatment of candidaemia and invasive candidiasis [16]. Whether combinations of isavuconazole with echinocandins possess synergistic interactions against *C. auris*, especially against biofilms, has been poorly studied. Hence, we examined *in vitro* and *in vivo* combinations of isavuconazole and caspofungin against *C. auris* isolates derived from the four main clades.

## 2. Material and Methods

### 2.1. Isolates

Isolates of four different *C. auris* clades (South Asian, n = 9; East Asian, n = 4; South African, n = 5; South American, n = 5) were tested; their origin is listed in Supplementary Table 1. All isolates were identified to the species level by matrix-assisted laser desorption/ionisation time-of-flight mass spectrometry. Clade delineation was conducted by polymerase chain reaction (PCR) amplification and sequencing of the 28S ribosomal DNA (rDNA) gene and the internal transcribed spacer region 1, as described previously [17–18].

### 2.2. Determination of the planktonic minimal inhibitory concentration

The planktonic MIC (pMIC) was determined according to the recommendations proposed by the Clinical Laboratory Standards Institute M27-A3 protocol [19]. Susceptibility to caspofungin pure powder (Molcan, Toronto, Canada) and isavuconazole pure powder (Merck, Budapest, Hungary) was determined in RPMI-1640 (with L-glutamine and without bicarbonate, pH 7.0, and with MOPS; Merck, Budapest, Hungary). The drug concentrations tested ranged from 0.008 to 4 mg/L for isavuconazole and from 0.03 to 2 mg/L for caspofungin. pMICs were determined as the lowest drug concentration that produces at least 50% growth reduction compared with the growth control. pMICs represent three independent experiments per isolate and are expressed as the median. *Candida parapsilosis* ATCC 22019 and *Candida krusei* ATCC 6258 were used as quality control strains.

### 2.3. Biofilm development

*C. auris* isolates were subcultured on Sabouraud dextrose agar (Lab M Ltd., Bury, United Kingdom). After 48 hours, fungal cells were harvested by centrifugation (3000 *g* for 5 min) and were washed three times in sterile physiological saline. After the final washing step, pellets were resuspended in physiological saline (ca. 5–6 mL) and were counted using a Bürker chamber (Hirschmann Laborgera□te GmbH & Co. KG, Eberstadt, Germany). The final density of inoculums was adjusted in RPMI-1640 broth to 1 × 10^6^ cells/mL and 100 μL aliquots were inoculated onto flat-bottom 96-well sterile microtitre plates (TPP, Trasadingen, Switzerland) and then incubated statically at 37°C for 24 hours to produce one-day-old biofilms [20–22].

### 2.4. Determination of the minimal inhibitory concentration of one-day-old biofilms

The examined caspofungin concentrations for sessile MIC (sMIC) determination ranged from 1 to 32 mg/L, while the examined isavuconazole concentrations ranged from 0.008 to 2 mg/L. One-day-old biofilms were washed three times with sterile physiological saline. Subsequently, sMICs were determined in RPMI-1640 using a metabolic activity change–based XTT assay. The percentage change in metabolic activity was calculated based on absorbance (*A*) at 492 nm as 100% × (*A*_well_ – *A*_background_)/(*A*_drug-free well_ – *A*_background_). sMICs were defined as the lowest drug concentration resulting in at least a 50% metabolic activity decrease compared with untreated control cells [20–22]. sMICs represent three independent experiments per isolate and are expressed as the median.

### 2.5. Evaluation of interactions by fractional inhibitory concentration index and the Bliss independence model

Interactions between caspofungin and isavuconazole were assessed by a two-dimensional broth microdilution chequerboard assay [20–22]. Interactions were then analysed by determining the fractional inhibitory concentration index (FICI) and using the Bliss independence model [14, 20–23]. In the case of planktonic cells, the tested concentration ranged from 0.008 to 2 mg/L for isavuconazole and from 0.015 to 1 mg/L for caspofungin. For biofilms, the examined caspofungin concentrations ranged from 1 to 32 mg/L, while the tested isavuconazole concentrations ranged from 0.008 to 2 mg/L. FICIs were calculated with the widely used following formula: ΣFIC = FIC_A_ + FIC_B_ = [(MIC_A_^comb^/MIC_A_^alone^)] + |(MIC_B_^comb^/ MIC_B_^alone^)], where MIC_A_^alone^ and MIC_B_^alone^ stand for MICs of drugs A and B when used alone, and MIC_A_^comb^ and MIC_B_^comb^ represent the MICs of drugs A and B in combination at isoeffective combinations, respectively [14, 20–23]. FICIs were determined as the lowest ΣFIC. MICs of the drugs alone and of all isoeffective combinations were determined as the lowest concentration resulting in at least 50% metabolic activity reduction compared with the untreated control biofilms. If the obtained MIC was higher than the highest tested drug concentration, the next highest twofold concentration was considered the MIC. FICIs ≤ 0.5 were defined as synergistic, between > 0.5 and 4 as indifferent, and > 4 as antagonistic [14, 20–23]. FICIs were determined in three independent experiments and are presented as the median.

To further evaluate caspofungin–isavuconazole interactions, MacSynergy II analysis was applied; this approach employs the Bliss independence algorithm in a Microsoft Excel–based interface to determine synergy [20–24]. The Bliss independence algorithm is a well-described method for the examination of the nature of drug–drug interactions. Briefly, the Bliss independence algorithm calculates the difference (ΔE) in the predicted percentage of growth (E_ind_) and the experimentally observed percentage of growth (E_exp_) to define the interaction of the drugs used in combination. E_ind_ is calculated with the equation E_ind_ = E_A_ × E_B_, where E_ind_ is the predicted percentage of growth that defines the effect of combination when the drugs are acting alone. E_A_ and E_B_ are the experimental percentages of growth with each drug acting alone. The MacSynergy II model uses interaction volumes and defines positive volumes as synergistic and negative volumes as antagonistic. The obtained E values of each combination are presented on the z-axis in the three-dimensional plot. Synergy or antagonism is significant if the interaction log volumes are higher than 2 or lower than 2, respectively [14, 24]. Log volume values between > 2 and 5, between > 5 and 9, and > 9 should be considered as minor synergy, moderate synergy and strong synergy, respectively. The corresponding negative values define minor, moderate and strong antagonism, respectively. The synergy volumes were calculated at the 95% confidence level [24].

### 2.6. Infection model

Pathogen-free female BALB/c mice weighing 22 to 24 g were used for the *in vivo* experiments. The Guidelines for the Care and Use of Laboratory Animals were strictly followed during the maintenance of mice. Animals were allowed access to food and water *ad libitum. In vivo* experiments were approved by the Animal Care Committee of the University of Debrecen (permission number is 12/2014). An immunocompromised mouse disseminated model was used for the studies. The animals were rendered neutropenic by intraperitoneal injection of cyclophosphamide (Endoxan, Baxter, Deerfield, IL, United States) 4 days (150 mg/kg body weight) and 1 day (100 mg/kg body weight) before infection and then 2 and 4 days postinfection (100 mg/kg body weight) [25]. Mice were infected intravenously through the lateral tail vein with 1–1.3 × 10^7^ colony-forming units (CFU) in 200 μL physiological saline [25]. The inoculum density was confirmed by plating serial dilutions on Sabouraud dextrose agar. Mice were divided into four groups (8 mice per group): (i) untreated control; (ii) 1 mg/kg/day caspofungin; (iii) 20 mg/kg/day isavuconazole; and (iv) 1 mg/kg/day caspofungin + 20 mg/kg/day isavuconazole. Cresemba© intravenous formulation (Basilea Pharmaceutica Ltd., Basel, Switzerland) was used for isavuconazole treatment. All treatment arms were given intraperitoneally and started 24 hours postinfection. In the case of caspofungin–isavuconazole combination, isavuconazole doses were administered 1 hour after the caspofungin treatments. Control mice were given 0.5 mL sterile physiological saline intraperitoneally. At 6 days postinfection, animals were euthanised; subsequently, the kidneys of each mouse were removed, weighed and homogenised aseptically. Homogenates were serially diluted tenfold and 100 μL aliquots were plated onto Sabouraud dextrose agar for viable fungal colony counts after incubation for 48 hours at 37°C [25]. The lower limit of detection was 500 CFU/kidney. The kidney burden was analysed using the Kruskal–Wallis test with Dunn’s post-test (GraphPad Prism 6.05.). Significance was defined as p < 0.05.

## 3. Results

The median and the range of MICs for planktonic and sessile *C. auris* isolates are shown in Table 1. The planktonic form of the tested isolates was considered to be susceptible to caspofungin based on the tentative MIC breakpoint recommended by the Centers for Disease Control and Prevention (≥ 2 mg/L) [26]. By the microdilution method, the 23 isolates exhibited pMICs for caspofungin alone from 0.5 to 2 mg/L, with a pMIC_50_, pMIC_90_ and geometric mean pMIC of 1, 2 and 1.13 mg/L, respectively. In the case of isavuconazole, pMICs were from 0.015 to 4 mg/L, with a pMIC_50_, pMIC_90_ and geometric mean pMIC of 0.5, 2 and 0.33 mg/L, respectively. Fourteen out of 23 isolates formed biofilms, which showed significantly higher resistance to caspofungin and isavuconazole compared with planktonic cells (Table 1). sMICs for caspofungin alone were from 2 to > 32 mg/L, with a sMIC_50_, sMIC_90_ and geometric mean sMIC of > 32, > 32 and 45.25 mg/L, respectively (64 mg/L was used for geometric mean sMIC analysis in the case of sMIC > 32 mg/L). The biofilm-forming isolates exhibited sMICs for isavuconazole alone from 0.5 to > 2 mg/L, with a sMIC_50_, sMIC_90_ and geometric mean sMIC of > 2, > 2 and 3.12 mg/L, respectively (4 mg/L was used for geometric mean sMIC analysis in the case of sMIC > 2 mg/L) (Table 1).

**Table 1.** Minimum inhibitory concentrations (MICs) of caspofungin alone and in combination with isavuconazole against *Candida auris* planktonic cells and one-day-old biofilms.

| Clades | Isolates | Planktonic cells<br>Median MIC (range) of drug used (50% O.D. reduction in turbidity) |  |  |  | Biofilms<br>Median MIC (range) of drug used (50% O.D. reduction in metabolic activity) |  |  |  |
| --- | --- | --- | --- | --- | --- | --- | --- | --- | --- |
|  |  | Alone |  | In combination |  | Alone |  | In combination |  |
|  |  | Caspofungin (mg/L) | Isavuconazole (mg/L) | Caspofungin (mg/L) | Isavuconazole (mg/L) | Caspofungin (mg/L) | Isavuconazole (mg/L) | Caspofungin (mg/L) | Isavuconazole (mg/L) |
| South Asian | 10 | >1<sup>a</sup> | >2<sup>b</sup> (2- >2) | 0.25 (0.25-0.5) | 0.015 (0.008-0.015) | >32<sup>c</sup> | >2<sup>b</sup> | 0.5 | 0.015 (0.015-0.06) |
|  | 12 | 1 | 1 (0.5-1) | 0.5 (0.015-0.5) | 0.015 (0.015-0.25) | >32<sup>c</sup> | >2<sup>b</sup> | 0.5 | 0.015 (0.008-0.015) |
|  | 20 | >1<sup>a</sup> | 2 (2- >2) | 1 | 0.015 (0.015-1) | >32<sup>c</sup> | >2<sup>b</sup> | 0.5 | 0.03 |
|  | 27 | >1<sup>a</sup> (1- >1) | 1 (0.5- >2) | 0.25 (0.25-0.5) | 0.015 (0.015-0.06) | >32<sup>c</sup> | >2<sup>b</sup> | 0.5 | 0.015 (0.008-0.015) |
|  | 33 | >1<sup>a</sup> | 0.06 | 0.015 (0.015-0.03) | 0.008 | NA | NA | NA | NA |
|  | 82 | >1<sup>a</sup> | 2 | 0.25 (0.25-1) | 0.25 (0.03-0.25) | >32<sup>c</sup> | >2<sup>b</sup> | 4 (4-8) | 0.06 (0.008-0.06) |
|  | 164 | >1<sup>a</sup> | 0.5 | 0.5 (0.5-1) | 0.06 (0.03-0.06) | NA | NA | NA | NA |
|  | 174 | >1<sup>a</sup> | 0.06 | 0.015 (0.015-0.25) | 0.008 | NA | NA | NA | NA |
|  | 196 | >1<sup>a</sup> | 0.06 | 0.06 (0.06-0.125) | 0.008 (0.008-0.015) | NA | NA | NA | NA |
| East Asian | 15 | 0.5 | 0.5 (0.25-0.5) | 0.25 | 0.25 | >32<sup>c</sup> | >2<sup>b</sup> | 0.5 | 0.125 |
|  | 12372 | 1 | 0.5 | 0.125 | 0.008 (0.008-0.015) | >32<sup>c</sup> | >2<sup>b</sup> | 4 | 1 |
|  | 12373 | 0.5 | 0.5 | 0.25 | 0.25 (0.25-0.125) | NA | NA | NA | NA |
|  | Type strain (NCPF 13029) | 0.5 | 0.03 | 0.015 | 0.008 | NA | NA | NA | NA |
| South African | 2 | 1 | 0.125 (0.06-0.125) | 0.015 | 0.008 (0.008-0.015) | NA | NA | NA | NA |
|  | 185 | 1 | 0.25 (0.125-0.25) | 0.015 | 0.03 (0.03-0.015) | NA | NA | NA | NA |
|  | 204 | 0.5 | 0.125 (0.06-0.125) | 0.015 | 0.03 | 32 (16-32) | >2<sup>b</sup> | 0.5 | 0.008 |
|  | 206 | 1 | 0.25 (0.125-0.25) | 0.015 | 0.008 | NA | NA | NA | NA |
|  | 228 | 1 | 0.06 | 0.03 (0.015-0.06) | 0.008 (0.008-0.015) | 32 | >2<sup>b</sup> | 0.5 | 0.008 (0.008-0.015) |
| South American | I-24 | >1<sup>a</sup> | 1 (0.5-1) | 0.015 (0.015-1) | 0.008 (0.008-0.03) | >32<sup>c</sup> | 0.5 | 4 | 0.125 |
|  | I-172 | 1 | 0.5 | 0.5 | 0.25 | >32<sup>c</sup> | 0.5 (0.5-2) | 16 | 0.06 |
|  | 13108 | >1<sup>a</sup> | 2 | 1 | 0.008 | >32<sup>c</sup> | >2<sup>b</sup> | >32<sup>c</sup> | >2<sup>b</sup> |
|  | 13112 | 0.5 | 2 | 0.015 (0.015-0.03) | 0.5 | >32<sup>c</sup> | >2<sup>b</sup> | >32<sup>c</sup> | >2<sup>b</sup> |
|  | 16565 | 0.5 | 0.015 | 0.015 | 0.008 | 2 | 1 (0.5-1) | 0.5 | 0.06 |

<sup>a</sup>MIC is off-scale at >1 mg/L, 2 mg/L (one dilution higher than the highest tested concentration) was used for FICI analysis.
<sup>b</sup>MIC is off-scale at >2 mg/L, 4 mg/L (one dilution higher than the highest tested concentration) was used for FICI analysis.
<sup>c</sup>MIC is off-scale at >32 mg/L, 64 mg/L (one dilution higher than the highest tested concentration) was used for FICI analysis.

The median pMICs observed in combination showed a 2–128-fold and a 2–256-fold reduction for caspofungin and isavuconazole, respectively. A similar marked reduction in median sMICs was observed for biofilms (a 0–128-fold and a 0–512-fold decrease for caspofungin and isavuconazole, respectively) (Table 1).

Table 2 summarises the *in vitro* interactions between caspofungin and isavuconazole based on the median FICIs. An antagonistic interaction was never observed (all FICIs ≤ 4). Using a two-dimensional broth microdilution chequerboard assay and FICI calculation, the nature of the caspofungin–isavuconazole interaction was synergistic for 61% of the planktonic isolates, with median FICIs from 0.03 to 0.5 and a mean of the median FICI of 0.34. In the case of sessile cells, synergism was observed for 86% of the 14 biofilm-forming isolates, with median FICIs from 0.029 to 0.5 and a mean of the median FICI of 0.14 (Table 2).

**Table 2.** *In vitro* interactions by fractional inhibitory concentration indices (FICIs) of caspofungin in combination with isavuconazole against *Candida auris* planktonic cells and one-day-old biofilms.

| Clades | Isolate | Planktonic cells |  | Biofilms |  |
| --- | --- | --- | --- | --- | --- |
|  |  | FICI |  | FICI |  |
|  |  | Median (range) of FICI | Interaction | Median (range) of FICI | Interaction |
| South Asian | 10 | 0.28 (0.28-0.312) | Synergy | 0.037 (0.037-0.068) | Synergy |
|  | 12 | 0.03 (0.03-1) | Synergy | 0.076 (0.023-0.076) | Synergy |
|  | 20 | 0.51 (0.51-0.75) | Indifferent | 0.023 | Synergy |
|  | 27 | 0.5 (0.25-0.5) | Synergy | 0.029 (0.029-0.038) | Synergy |
|  | 33 | 0.313 (0.258-0.375) | Synergy | NA | NA |
|  | 82 | 0.375 (0.25-0.51) | Synergy | 0.155 (0.133-0.155) | Synergy |
|  | 164 | 0.62 (0.56-0.75) | Indifferent | NA | NA |
|  | 174 | 0.383 (0.375-0.5) | Synergy | NA | NA |
|  | 196 | 0.53 (0.5-0.625) | Indifferent | NA | NA |
| East Asian | 15 | 1 (0.75-1) | Indifferent | 0.5 | Synergy |
|  | 12372 | 0.37 (0.31-0.5) | Synergy | 0.375 (0.187-0.375) | Synergy |
|  | 12373 | 0.5 (0.5-1) | Synergy | NA | NA |
|  | Type strain (NCPF 13029) | 0.296 | Synergy | NA | NA |
| South African | 2 | 0.37 (0.255-0.49) | Synergy | NA | NA |
|  | 185 | 0.245 (0.245-0.255) | Synergy | NA | NA |
|  | 204 | 0.51 (0.49-0.54) | Indifferent | 0.019 | Synergy |
|  | 206 | 0.255 (0.255-0.3) | Synergy | NA | NA |
|  | 228 | 0.53 (0.5-0.625) | Indifferent | 0.03 (0.03-0.06) | Synergy |
| South American | I-24 | 0.51 | Indifferent | 0.5 (0.5-0.56) | Synergy |
|  | I-172 | 1 | Indifferent | 0.31 (0.28-0.37) | Synergy |
|  | 13108 | 0.5 | Synergy | 2 | Indifferent |
|  | 13112 | 0.37 (0.37-0.5) | Synergy | 2 | Indifferent |
|  | 16565 | 0.563 (0.563-0.593) | Indifferent | 0.31 | Synergy |

FICI calculation involves Loewe additivity-based analysis assuming that both drugs have the same mechanism of action, while the Bliss independence–based MacSynergy II program does not have this assumption. Figure 1 shows the dose-response surfaces for caspofungin– isavuconazole generated with MacSynergy II. Based on clade-specific cumulative log volumes, the combination of caspofungin and isavuconazole exerted minor synergy (the synergy log volume was 4.83) against planktonic isolates derived from the South African clade (Figure 1C). For the South Asian, South American and East Asian clades, the synergy log volumes were zero, indicating indifferent interactions (Figure 1A, B and D). In the case of biofilms, 77.2, 23.21 and 234.32 cumulative synergy log volumes were observed for South African, South Asian and East Asian clades, respectively, indicating strong synergistic interactions (Figure 1E, G and H). By contrast, the South American clade exhibited an indifferent interaction, with a cumulative synergy log volume of 0.13 (Figure 1F). Based on the evaluation of *in vitro* combinations, the data derived from the FICI calculation correlate with the MacSynergy analysis primarily in the case of biofilms. Although the combination of caspofungin and isavuconazole was synergistic or considerably reduced the amount of drug needed in some instances, the observed results may show strain specificity within clades, especially in the case of planktonic cells.

**Figure 1.**
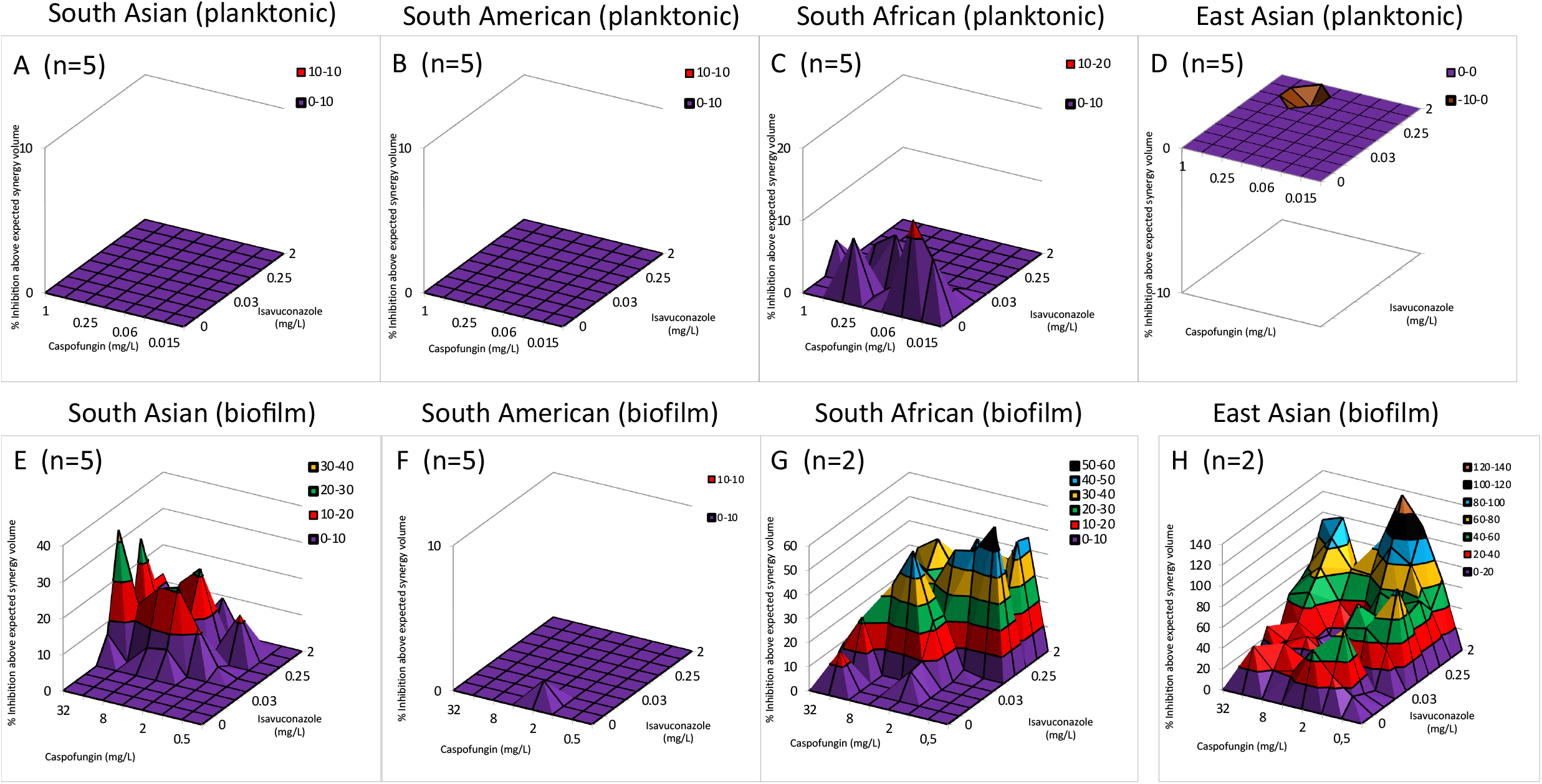
Effect of caspofungin in combination with isavuconazole against planktonic (A-D) and sessile (E-H) *Candida auris* isolates using MacSynergy II analysis. Positive values show synergy, while negative values indicate antagonism at given concentrations. The volumes are calculated at the 95% confidence interval.

To further evaluate the *in vivo* applicability of the caspofungin and isavuconazole combination, representative isolates were chosen where synergistic and indifferent interactions were observed *in vitro*, respectively. The results of the *in vivo* experiments are shown in Figure 2. One mg/kg daily caspofungin treatment decreased the fungal kidney burden in the case of the tested isolates; however, this therapeutic strategy was not statistically different compared with untreated control mice (p > 0.05). The 20 mg/kg/day isavuconazole treatment proved to be statistically ineffective against the tested *C. auris* isolates, especially in the case of isolate 13112 (p > 0.05). It is noteworthy that the fungal tissue burden decreases were higher than the three log decreases in mice treated with a daily combination of 1 mg/kg caspofungin and 20 mg/kg isavuconazole, which was statistically significant compare with control mice (p < 0.001) (Figure 2).

**Figure 2.**
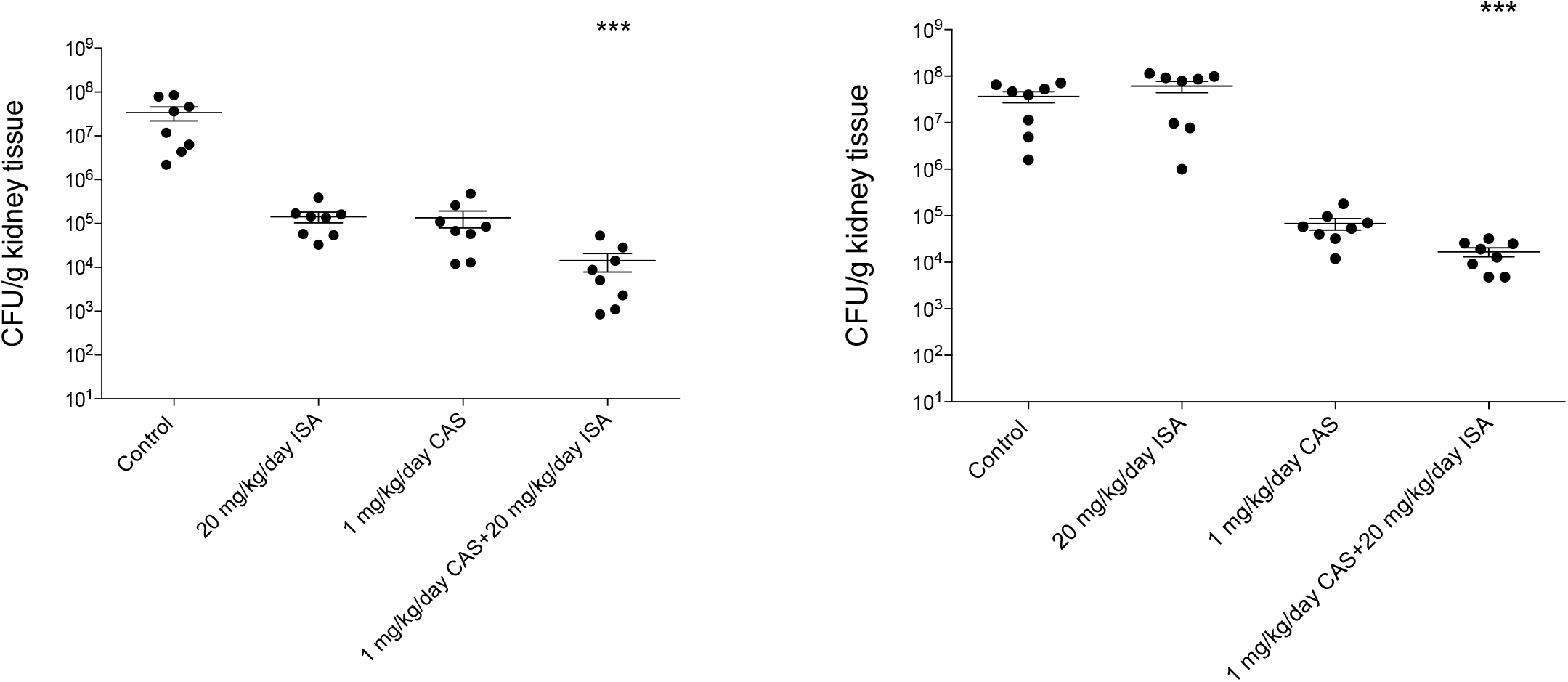
Kidney tissue burden of deeply neutropenic BALB/c mice infected intravenously with *Candida auris* 12 (A) and 13112 (B) isolates. Daily intraperitoneal caspofungin (CAS) (1mg/kg/day) and isavuconazole (ISA) (20 mg/kg/day) treatment was started 24 hours after the infection. Tissue burden experiments were performed on day 6 post-infection. Bars represent means ± standard error of mean. *** corresponds to p<0.001 compared with the control population.

## 4. Discussion

The impending challenge of antifungal resistance and newly emerged fungal pathogens necessitates bold and innovative therapeutic solutions [27]. In recent years, combination-based antifungal treatments have become a promising therapeutic approach, especially against multidrug-resistant fungal species such as *C. auris*. Based on previous susceptibility studies against *C. auris*, the efficacy of *in vitro* combinations has shown high variability; in addition, the degree of activity is highly strain – or rather clade – specific [28–30].

Isavuconazole is recommended primarily for the treatment of invasive aspergillosis and mucormycosis; however, it has also exerted variable *in vitro* activity against several *Candida* species [15]. Sanglard and Coste (2016) reported that the activity range of isavuconazole is similar to that of voriconazole against the *Candida* strains they tested, findings that were confirmed by Marcos-Zambrano *et al*. (2018), who showed high *in vitro* activity of isavuconazole against clinically relevant *Candida* species, particularly against *C. albicans* [31, 32]. Desnos-Ollivier *et al*. (2019) reported an isavuconazole MIC of 0.015 mg/L against planktonic *C. auris*; however, they tested only two strains [33]. Regarding clinical findings, the ACTIVE trial compared intravenous isavuconazole to intravenous caspofungin followed by oral isavuconazole in a phase 3 randomised, double-blind clinical trial for patients with *Candida* bloodstream infection. These results support the use of isavuconazole as a potential therapy for candidiasis [16].

To the best of our knowledge, this is the first study to examine the *in vitro* and *in vivo* combined effect of isavuconazole and caspofungin against *C. auris* strains derived from four different lineages focusing on both planktonic and sessile susceptibility. In the case of *Aspergillus* spp., this combination showed synergistic interaction in 13% of tested strains [34]. In our study, we found *in vitro* synergy for the caspofungin–isavuconazole combination using chequerboard microdilution, especially based on FICI determination, which was definitely pronounced in the case of one-day-old biofilms. Katragkou *et al*. [35] showed synergistic interactions between isavuconazole and micafungin against *C. albicans, Candida parapsilosis* and *Candida krusei* using the Bliss independence model (the degree of synergy ranged from 1.8% to 16.7%), which was confirmed by time-kill curves, especially against *C. albicans* and *C. parapsilosis*. Voriconazole exerted synergistic interaction with caspofungin or other echinocandins against *C. auris* isolates using the FICI [36]. In a recent study, Pfaller *et al*. [37] examined the *in vitro* activity of voriconazole or isavuconazole in combination with anidulafungin; synergy or partial synergy was observed in 14% and 61% of the isolates with the combination of anidulafungin plus voriconazole and in 19% and 53% of isolates for the combination of anidulafungin plus isavuconazole. It is noteworthy that O’Brien *et al*. [29] examined four pan-resistant *C. auris* strains derived form a New York outbreak to evaluate whether they are susceptible to combinations of antifungals. Based on their results, flucytosine combinations with either amphotericin B, azoles or echinocandins exhibited the highest efficacy [29]. However, the combination of azoles with echinocandins had no superior effect compared with monotherapy [29].

The number of *in vivo* experiments focusing on combination-based therapy against *C. auris* is strongly limited. In the only published *in vivo* combination-based experiments, Eldesouky *et al*. [38] observed that the examined sulfamethoxazole–voriconazole combination enhanced the survival of *Caenorhabditis elegans* nematodes infected with *C. auris* by nearly 70%. Our study is the first that has examined the effect of caspofungin in combination with isavuconazole *in vivo* at clinically relevant concentrations using an immunocompromised mouse model. Although caspofungin alone produced a remarkable reduction in the kidney fungal burden, only its combination with isavuconazole was statistically superior compared with the untreated control (p < 0.001).

The multidrug resistance phenotype is a well-known characteristic for *C. auris*; it may be more pronounced in biofilms and further complicate treatment [6]. Based on previous susceptibility testing, amphotericin B, fluconazole, voriconazole, anidulafungin, micafungin and caspofungin could not completely eradicate *C. auris* biofilms *in vitro*, increasing the need for effective combination therapies [39]. Certain non-antifungal agents in combination with traditional antifungal drugs have been tested to eradicate *C. auris* biofilms with variable efficacy [21, 22, 40]. However, to date there is no experimental evidence about the efficacy of antifungal drug–drug combinations against *C. auris* biofilms. In our study, we found a prominent synergistic interaction between caspofungin and isavuconazole against biofilms for three out of the four clades examined. We observed indifferent interaction only in the case of two hospital-derived isolates from the South American clade (13112, 13108). The origin of these strains may explain the significantly higher resistance against drugs tested and the observed indifferent interaction compared to other isolates.

It should be pointed out that our study had a limitation, namely the choice of the endpoint for FICI-based assessment of antifungal combinations. To date, there is no a solid consensus about which endpoint should be used [23, 24, 30]. In addition, for MacSynergy-based evaluation, there is no endpoint at all, and the nature of the interaction is calculated only based on the percentage of growth at given concentrations [24, 30]. Despite this limitation, the therapeutic potential of the caspofungin and isavuconazole in combinations is unquestionable, which was definitely confirmed against *C. auris* biofilms and our *in vivo* experiments.

In summary, the presented synergistic combinations correspond to clinically achievable and safe drug concentrations. Our findings suggest that administration of the caspofungin– isavuconazole combination may help to expand the therapeutic options against *C. auris*. Nevertheless, the more extensive *in vivo* correlation and significant clinical relevance of these *in vitro* and *in vivo* results warrants further studies, especially in the case of biofilms.

## Ethical approval

Not required.

