## Supplemental table 1 for "*In vitro* and *in vivo* interaction of caspofungin with isavuconazole against *Candida auris* planktonic cells and biofilms": Supplementary table 1_.pdf

**Supplementary table 1** Origin of *Candida auris* isolates used in the study

| Clade | Isolate number | Country | Body site |
|---|---|---|---|
| **South Asian** | **10** | England | Wound swab |
| | **12** | England | Not stated |
| | **20** | England | Wound swab |
| | **27** | England | Pleural fluid |
| | **33** | England | Blood |
| | **82** | England | Blood |
| | **164** | England | Swab |
| | **174** | England | Nose |
| | **196** | Oman | Blood |
| **East Asian** | **15** | England | Not stated |
| | **12372** | Korea | Blood |
| | **12373** | Korea | Blood |
| | **Type strain (NCPF 13029)** | Japan | External ear |
| **South African** | **2** | England | Cerebrospinal fluid |
| | **185** | England | Blood |
| | **204** | England | Tracheostomy |
| | **206** | England | Blood |
| | **228** | England | Skin swab |
| **South American** | **I-24** | Israel | Blood |
| | **I-172** | Israel | Blood |
| | **13108** | Panama | Hospital environment |
| | **13112** | Panama | Hospital environment |
| | **16565** | Chicago (from Colombia) | Hospital environment |
